# Provirus induction diversifies adaptive variation in *Pseudomonas aeruginosa* lysogen populations

**DOI:** 10.64898/2026.01.04.697566

**Authors:** Laura C. Suttenfield, Mia Mercado, Sanjana Jose, Ahmad Hassan, Rachel J. Whitaker

**Affiliations:** Department of Microbiology, School of Molecular and Cellular Biology, University of Illinois Urbana-Champaign, Urbana, IL; Carl R. Woese Institute for Genomic Biology, University of Illinois Urbana-Champaign, Urbana, IL; Department of Biomedical Informatics, Harvard Medical School, Boston, MA

**Keywords:** Lysogeny, phage, provirus, bacteria, evolution, clonal interference, adaptive mutation, antibiotic resistance, *Pseudomonas aeruginosa*, Mu-like, DMS3

## Abstract

*Pseudomonas aeruginosa* is a Gram-negative opportunistic pathogen that forms chronic infections in people with cystic fibrosis. Often *P. aeruginosa* strains are lysogens, infected with proviruses, that are induced by cellular stressors which include challenge with antibiotics such as ciprofloxacin. We asked how infection of *P. aeruginosa* with Mu-like provirus DMS3 would alter the evolution of antibiotic resistance through serial transfer of six replicates of four strain backgrounds, with and without sublethal doses of ciprofloxacin over 20 days. Through population metagenome sequencing we found that inducing lysogen populations had significantly higher diversity in adaptive mutations than non-lysogens after 20 days. Lysogen populations have more co-existing mutations within the population and a higher number of mutations in adaptive genes, including antibiotic efflux pump and gyrase alleles, and CRISPR-Cas loci. These data suggest that viral induction shifts the adaptive regime of a clonal population from one of stepwise periodic selective sweeps to recurrent competing mutations. Our results demonstrate the way that provirus infection can shape the evolutionary trajectory of their hosts in an environment-dependent manner.

**Significance:** Most long-term chronic infections of *Pseudomonas aeruginosa* in cystic fibrosis patients trace their origin to a single ancestor. During long-term chronic infection the evolution of each unique ancestor strain, defined by its starting genotype, can dramatically impact the success of future treatment efforts. Often the common ancestral *Pseudomonas aeruginosa* genotype is infected by a virus; this in turn shapes the evolution of bacterial cells within the human island of the lung. This work shows that when the nested virus infecting *Pseudomonas aeruginosa* is the very common transposable Mu-like virus, it can cause the bacterial population to diversify in response to antibiotic treatment in ways that may complicate future antibiotic treatment and phage therapy.

## Introduction

The majority of sequenced bacteria, are lysogens, infected with one or more temperate viruses (proviruses) (*1–5*). Lysogeny can reconfigure host physiology (*6*) and be a source of evolutionary innovation or conflict between the virus genome and the bacterial chromosome (*7–9*). Lytic activation of the provirus (known as induction) can occur spontaneously or be triggered by a variety of stresses, such as nutrient limitation, chemical stress, or DNA damage (*10*). Induction of transposable proviruses results in viral transposition around the genome, increasing rates of mutation as the virus replicates through repeated insertion in the chromosome (*11*). Provirus induction can alter demography, growth rate, mutation rate and the landscape of adaptive mutations in lysogen populations. Therefore, proviruses alter the evolutionary trajectory of bacterial chromosome and may cause differences in the process of adaptative evolution of lysogen populations compared to uninfected cells (*12*). Despite this, there have not yet been studies directly comparing lysogen and non-lysogen evolution under inducing conditions.

*Pseudomonas aeruginosa* has been widely studied as a model system for studying evolutionary dynamics in long-term adaptation to the lungs of patients with cystic fibrosis (CF), as infections often begin with low strain diversity and then expand clonally over the long-term chronic infection (*13*, *14*). Within a single patient, it is common that all strains trace their origin to a common ancestor; in the case of a lysogen, this results in an absence of susceptible hosts for the induced viruses to infect. Lysogeny is common in CF isolates and ∼20% of *P. aeruginosa* strains are infected by a large family of transposable, Mu-like viruses (*15*, *16*). One model for these Mu-like viruses, DMS3, was experimentally induced from *P. aeruginosa* isolated from a CF patient (*17*). The addition of free DMS3 and related viruses – including LES phi-4, a DMS3-like provirus which infects the transmissible *P. aeruginosa* Liverpool Epidemic strain (*18*) – to cultures of susceptible *P. aeruginosa* have been shown to contribute to variation in experimentally evolved populations through the creation of knockout mutations upon insertion (*12*, *19*). However, the forward evolution of an individual lysogen ancestor may be more relevant in environments like the CF lung, where clonal expansion of ancestral strains are common and spatial structure across lung compartments (*20*, *21*) and in biofilms (*22*) has been shown to favor temperate strategies in viruses (*23*) and decreased viral virulence even in purely lytic viruses (*24*).

Antibiotics such as ciprofloxacin, one of the few recommended therapies for CF patients infected with *P. aeruginosa* (*25*, *26*), can induce temperate proviruses (*16*, *27*, *28*). Ciprofloxacin binds to essential bacterial proteins DNA gyrase and topoisomerase IV, which both function to relieve supercoiling during replication by introducing and re-ligating double-stranded breaks (*29*). In the presence of ciprofloxacin or other fluoroquinolone antibiotics, unresolved double-stranded breaks are left in the chromosome that trigger viral induction and eventually cell death through viral lysis (*30*). Resistance is mediated through point mutations in DNA gyrase genes *gyrA* and *gyrB* and topoisomerase IV genes *parC* and *parE*, but may also arise via efflux pump overexpression (*31–36*).

Theoretical work based on *P. aeruginosa*–ciprofloxacin parameters predicts that proviral induction and lysis of *Pseudomonas aeruginosa* lysogens acts synergistically with ciprofloxacin treatment, requiring less antibiotic to achieve the same level of bacterial clearance (*37*). Experimentally, synergy between ciprofloxacin and temperate viruses of *P. aeruginosa* has been demonstrated, where the administration of temperate viruses increases bacterial sensitivity to ciprofloxacin even at sublethal concentrations through inducing newly formed lysogens (*38*). Previously, we showed that the DMS3 provirus infecting type strain PA14 contributed to genetic variation through active, within-genome transposition (*8*). Genetic conflict between PA14 and its provirus resulted in costly low levels of ongoing induction because of CRISPR self-targeting with a degraded CRISPR spacer match (*39*). After 12 days of evolution, we found that large deletions around this match mediated by virus transposition co-existed with strains with slightly higher levels of induction that had resolved genetic conflict through mutation (*8*).

Here we test how lysogens evolve resistance to ciprofloxacin in near-isogenic starting lysogen and nonlysogen populations to reflect homogeneous starting populations of *P. aeruginosa* in the CF lung. Our analysis shows that provirus induction shifts the evolutionary regime toward clonal interference and successive mutation resulting in a mutation-selection balance with higher standing variation than non-inducing populations in a rugged evolutionary landscape.

## Methods

### MIC of ancestral strains in liquid media

We confirmed that our strains are susceptible to ciprofloxacin by diluting mid-log phase cultures 100-fold into a 96-well deep well plate with ciprofloxacin concentrations which decreased 2-fold from 25 µg/mL. We grew these plates overnight and quantified PFUs through plaque assays and CFUs on LB agar plates without antibiotics from each well.

### Experimental evolution

Strains were evolved in 300 µL volumes in 1.2 mL deep well 96 well plates in LB, shaking at 220 RPM, and at 37°C. These strains differed in their CRISPR content: PA14_lys has a full CRISPR-cas locus and an imperfect match between a host spacer and the provirus, which causes low levels of DNA damage (*40*); ΔCRISPR_lys has a deletion of both CRISPR arrays and *cas* genes (*41*). PA14 and ΔCRISPR, our uninfected strains, differed only in their CRISPR content. The two lysogens have a virus insertion in the same locus (the intergenic space upstream of *phzM*, a gene in the pyocyanin biosynthetic pathway), but PA14_lys has an additional 20-kb deletion downstream of this site and does not produce pyocyanin.

Populations were inoculated from single colonies, in matching replicates; populations with the same number, of the same strain, are from the same colony. For example, “ΔCRISPR _CIP_2” and “ΔCRISPR _noCIP_2” populations both originate from the same ΔCRISPR colony. We transferred 1% of overnight culture to fresh media every 24 hours for 20 days. 0.05 μg/mL ciprofloxacin was added to +CIP treatments each day. Each day, we diluted the culture 1/10 and measured the OD in a plate reader; every three days, we spotted CFU and PFUs. To avoid contamination between wells, we evolved each strain in its own plate, stored lysogen and nonlysogen cultures for frozen stock in separate plates, and tested for PFU production from non-lysogen populations using spot-on-lawn assays every day.

### Growth curves and solid media MIC

To test the effect of ciprofloxacin on +CIP- and noCIP-evolved populations and their relative growth rates, we patched the evolved populations onto LB agar plates and inoculated 96 well plates from these patches by touching pipette tips to the patches and then the liquid media with a multichannel pipette. We inoculated all populations into 200 µL of LB with or without 0.05 µg/mL ciprofloxacin and incubated them in a plate reader with at 37°C for 24 hours. In order to measure antibiotic resistance in evolved strains, we grew up populations from overnight cultures begun from inoculating from evolved populations which were patched onto LB plates. We serially diluted these cultures and spotted onto solid LB agar with concentrations of ciprofloxacin decreasing 50% stepwise from 25 μg/mL and estimated CFUs from colony counts.

### Population sequencing

We extracted population gDNA from the frozen stock of Day 20 populations with a Qiagen DNeasy Blood and Tissue DNA extraction kit on a Biomek 4000 liquid handler (Beckman-Coulter). We sheared extracted population gDNA sonically with the Covaris M220 at a fragment length of 600 nt, and prepared libraries using Roche Kapa PCR-free library preparation to preserve relative abundance of allelic variants. Population DNA was pooled, and sequenced on a NovaSeq SP 2×250 lane by the Roy J. Carver Biotechnology Center at the University of Illinois Urbana-Champaign. Two populations were never inoculated (ΔCRISPR_noCIP_3 and ΔCRISPR_CIP_3), two were lost during passaging (PA14_CIP_5 and PA14_CIP_6) and one was a low-coverage outlier after sequencing and was subsequently excluded (ΔCRISPR_noCIP_5). In total, we analyzed 43 experimental and 4 ancestral control populations. 815 million reads in total were produced, with an average coverage of 397.4 ± 135.6.

### Genome assembly and analysis

We used an in-house pipeline to process the reads that included Trimmomatic to trim the reads and bwa-mem to align them to the reference (*42*). We used breseq (*43*) to identify mutations using polymorphism mode with default options, and samtools depth (*44*) to identify structural variations. Some repetitive regions had misassembly errors and unusually high numbers of mutations; we therefore masked all mutations from genes including PA14_40260, PA14_61200, and the intergenic space downstream of genes PA14_72510 and aceE. We used the Transposable Insertion Site Finder pipeline (available at github.com/igoh-illinois, (*8*)) to identify viral insertion sites, and confirmed these results by using the genome viewers Tablet 1.21.02.08 (*45*) and IGV 2.12.3 (*46*). We compared viral insertion site output from the “_hostSplits.tsv” output from the pipeline to our PA14 Genbank reference file with a custom script to assess new viral insertion site location, and divided read numbers by average coverage of the genome to approximate insertion site frequencies in the population. We manually annotated CRISPR deletions, and masked all genes that had viral insertions which formed a border of one of these deletions to avoid double-counting mutations. Finally, we filtered our dataset to include only insertions seen at >2% frequency in the population relative to each ancestral strain, and we subtracted all mutations that were also mutated in the ancestral metagenomes, except for mutations in *orfN*.

### Statistical analysis

All graphical and statistical analysis was done with R version 4.3.1 (*47*), using the packages ggplot v3.5.2 (*48*), dplyr v1.1.2 (*49*), vegan v2.6.8 (*50*), upsetR v1.4.0 (*51*) and rstatix v0.7.2 (*52*). Where necessary, we used log transformation in order to meet the assumptions for ANOVA, using the aov function from the stats package, v4.3.1. For post-hoc pairwise testing, we used the TukeyHSD function with our model object, also from the stats package. Area under the curve was calculated using the auc function from the flux package v0.3-0.1. Throughout, we used Kruskal-Wallis tests for pairwise comparisons with Bonferroni correction for multiple comparisons, and Kolmogorov-Smirnov (KS) tests to compare frequency distributions.

### Distance-based redundancy analysis

To ordinate Jaccard dissimilarities between populations and to decompose this dissimilarities into our variables, we constructed Jaccard similarity matrices on the presence/absence of the gene or mutation set, using the vegan::vegdist function with options “binary = TRUE” and “method = ‘jaccard’” in R (*47*, *50*). Significance of the model terms was calculated with the vegan::adonis2 function with the “by = ‘terms’” option, using the model format *matrix ∼ strain * cipro * lysogen:induction*. In order to control for plate location as a random effect, we set this variable as our strata in the how() function from the permute package (*53*) with 999 permutations, within = Within(type = “free”), plots = Plots(strata = *plate location*, type = “none”), and observed = true. Then, we used the dbrda function in the vegan package to create the ordination object in R. The vectors of the main terms are from the biplot output of the ordination object, while the vectors of individual genes, with their p-values, were extracted from the ordination object using the envfit function with the same permutations as in the adonis2 function.

### Data availability

All raw reads are available on NCBI under BioProject number XXXXX. Data and code to produce the figures are available at github.com/lsuttenfield/provirus-induction-evolution.

## Results

### Phenotypic variation

We used a GxGxE experimental evolution design, with strain background (+/- CRISPR), infection status (+/- DMS3 infection), ciprofloxacin addition (+/- sublethal Cip), and induction (inducing/noninducing) as variables. To establish the appropriate sublethal concentration of ciprofloxacin for our experimental conditions, and its impact on DMS3 induction in CRISPR self-targeting and ΔCRISPR DMS3 lysogens (PA14_lys and ΔCRISPR_lys, respectively), we challenged these lysogens with 0 – 0.4 µg/ml ciprofloxacin and measured CFUs and PFUs of overnight cultures (Figure S1A). The MIC of all strains was 0.2 µg/mL, as has been previously reported for *P. aeruginosa* (*54*), except PA14_lys, which was lower at 0.1 µg/mL due to DMS3 induction by CRISPR self-targeting. Sublethal concentrations of ciprofloxacin (0.05 µg/ml, 25% of the MIC for *P. aeruginosa,* Fig S1A) were sufficient to erase the difference in DMS3 plaque forming units (PFUs) between the CRISPR-containing and ΔCRISPR PA14 lysogens (Fig S1B, bottom right). This result suggests that sublethal ciprofloxacin is sufficient to induce to the level of CRISPR self-targeting and that CRISPR self-targeting and sublethal ciprofloxacin do not have noticeably additive effects on provirus induction. Therefore, we compared uninfected strains with and without CRISPR (PA14 and ΔCRISPR) to DMS3 lysogen strains with and without CRISPR (PA14_lys and ΔCRISPR_lys), and evolved them in parallel, in the presence and absence of sublethal ciprofloxacin (0.05 µg/mL) in six replicate populations each. We transferred 1% of the culture once every 24 hours for 20 days (∼6.7 generations/day, ∼134 generations total).

Figures 1A and 1B show the changes in bacterial growth of 43 populations (see Methods) as well as the average trajectory for each set of six replicates (Fig 1C). In all conditions that included sublethal ciprofloxacin evolution of resistance based on ODs at transfer (Fig 1B). The average OD after one day in ciprofloxacin was reduced for every strain compared to LB alone (Day 1 mean OD: 0.28 vs 1.55, CIP vs noCIP respectively). Over the course of the experiment, the OD increased ∼4-fold on average across all strains (Fig 1D, day 20 OD / day 1 OD = 3.94 ± 2.76). In noCIP-evolved populations, PFUs in self-targeting PA14_lys populations declined 10-fold over the 20 days, with a corresponding 4-fold increase of the terminal OD at transfer in PA14_lys (Fig 1A). In contrast, PFUs and OD were unchanged in the noninducing ΔCRISPR_lys populations.

**Figure 1.**
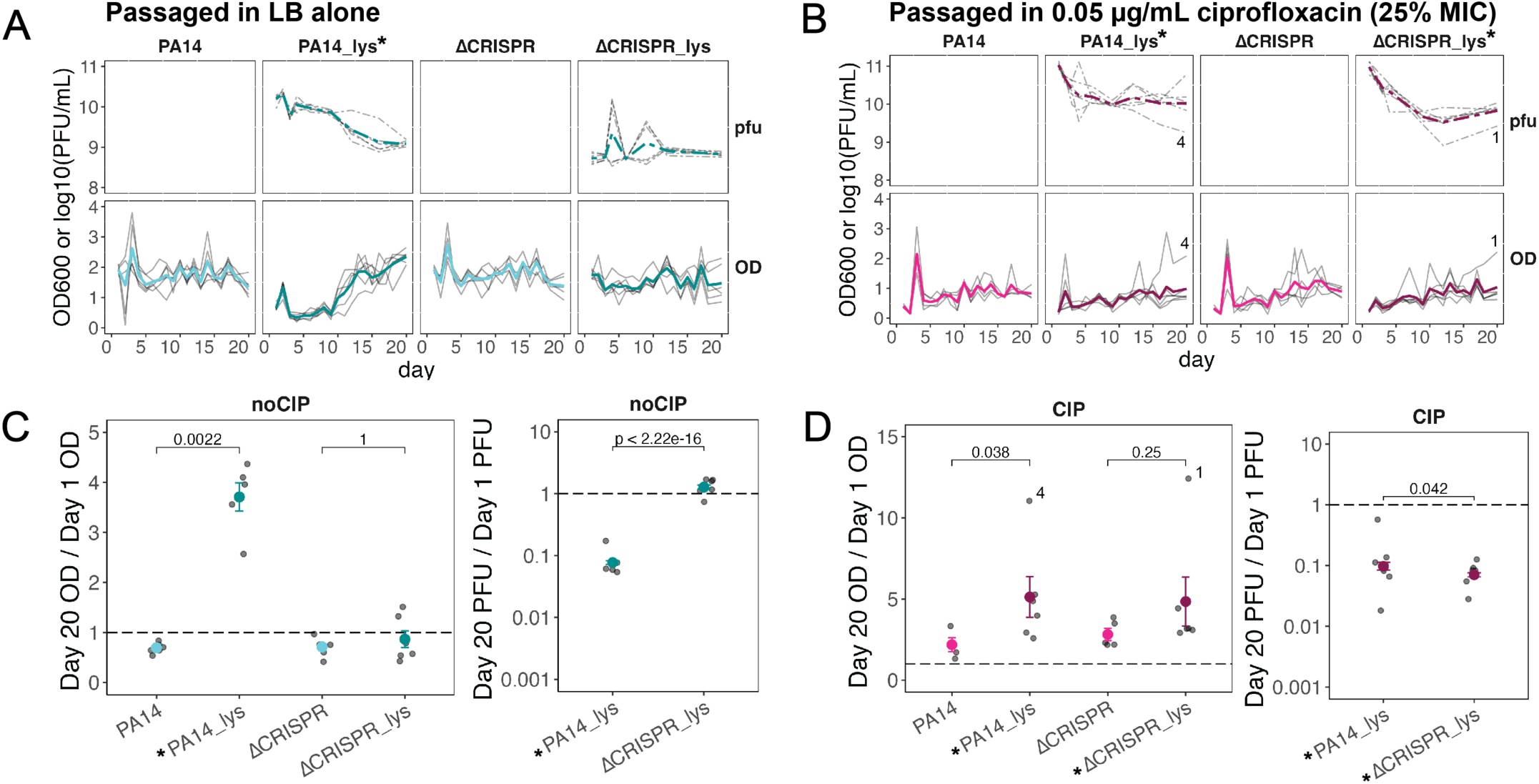
Lysogens resolve provirus induction from both CRISPR targeting and ciprofloxacin induction in culture. A, B) PFU/mL (top row, dashed lines) and OD600 (bottom row, solid lines) at the time of transfer, in A) LB alone media (noCIP) and B) media supplemented with 0.05 µg/mL ciprofloxacin (CIP). OD was taken every day and PFUs were quantified every three days. Numbers are population replicates. C, D) Ratios of Day 20 OD to Day 1 OD (left) or Day 20 PFU/mL over Day 1 PFU/mL (right) for each population evolved in noCIP and CIP. Points are the means; error bars are standard error. Significance of pairwise comparisons was calculated with Kruskal-Wallis tests. Asterisks denote strains that are inducing lysogens.

We assessed the level of resistance in each of the evolved populations on solid media with increasing concentrations of ciprofloxacin and saw that evolution in the presence of sublethal ciprofloxacin resulted in an average 23-fold increase in CIP resistance (Fig S1B; mean +CIP IC99 = 7 µg/mL, mean noCIP IC99 = 0.313 µg/mL). Comparing the IC99 of uninfected strains to lysogens in both backgrounds, differences between CIP-evolved lysogen and non-lysogen populations were not significant, and lysogen populations were significantly variable (Fig S1D). Whole-population growth curves in the presence and absence of sublethal ciprofloxacin showed that CIP-evolved inducing lysogen populations had slightly greater area under the curve (AUC) in the presence of ciprofloxacin (Fig S1E), but pairwise lysogen/nonlysogen comparisons were not significant. Phenotypically, lysogen and nonlysogen populations evolved equivalent levels of ciprofloxacin resistance.

PA14_lys noCIP, PA14lys CIP and ΔCRISPR_lys CIP had decreased PFUs and increased ODs at the end of the experiment, indicating adaptation to inducing conditions (Fig 1). In +CIP populations PFUs also decreased significantly in both lysogen backgrounds (Fig S2A), but stabilized at values an order of magnitude above those in noCIP, indicating that CIP continues to stimulate induction in CIP-resistant populations. There was higher variance among replicates in the +CIP evolved populations than existed in the noCIP condition; for example, PA14lys_CIP_4 and ΔCRISPRlys_CIP_1 had higher terminal OD600 values than other populations throughout the experiment, and lower PFUs (Fig 1B). Both populations harbored a fixed gyrase allele and as discussed in more detail below.

### Identifying adaptive mutations

To capture allelic diversity, we sequenced metagenomes of each of the six replicate cultures at the conclusion of the experiment using PCR-free library construction to avoid PCR bias in alleles frequencies (Methods). We used the combined outputs from *breseq* polymorphism mode (*43*) and a previously published tool for identification of transposable virus insertion sites (TISF, (*8*)), and compared them to our ancestral PA14, PA14_lys, ΔCRISPR and ΔCRISPR_lys reference metagenomes which were grown overnight.

We found 1,678 mutations (both nucleotide and transpositional) in 855 genes across all evolved populations which were present at frequencies ≥2% relative to our local reference PA14 strain (Fig 2A; Table 1). Nonsynonymous mutations were the largest category at 39% of the total (n = 659); synonymous mutations made up 13% of the total (n = 226). We recovered no mutations within the virus chromosome itself (Fig S2B); however, ciprofloxacin treatment significantly increased DMS3 coverage relative to the host chromosome in both lysogen backgrounds (Fig S2C), and we recovered a number of novel viral insertion sites across the genomes of these populations (Fig S2D).

**Figure 2.**
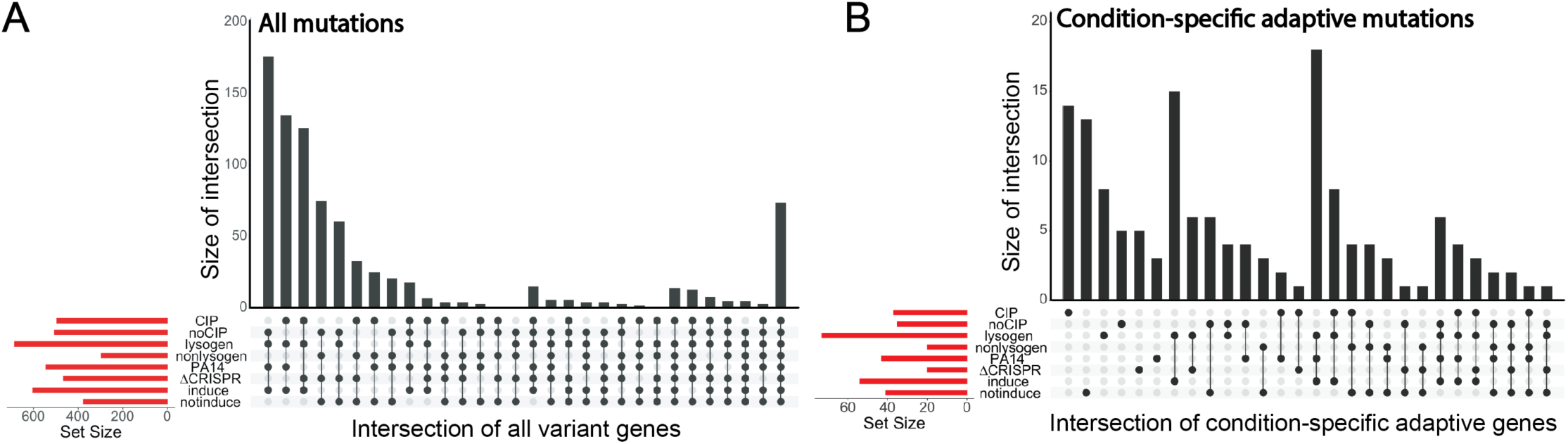
Shared and unique mutations across experimental conditions. A, B) Upset plot of A) all variant genes and B) condition-specific adaptive genes. Bars are the size of the intersection, which is represented as a grid black dots. The set size, or number of genes adaptive to each condition, is shown in the bar graph to the left.

**Table 1.** Counts of mutations in different categories across strain backgrounds, for both all and adaptive mutations. For both all mutations and adaptive mutations, counts shown are in terms of number of nucleotides and the percent of the total, summed across all replicate populations per strain x cipro background. Adaptive mutations are mutations in genes that are mutated in parallel across >1 population. Bold letters with asterisks indicate inducing strains. NS = nonsynonymous mutations; S = synonymous mutations; Ind = indels; Int = intergenic mutations; LgD = large deletions; Non = nonsense mutations; VI. = virus insertions.

| *Cipro* | *Strain* | *Mutation category* |  |  |  |  |  |  | Total | N: Genes |
| --- | --- | --- | --- | --- | --- | --- | --- | --- | --- | --- |
|  |  | NS | S | Ind | Int | LgD | Non | VI |  |  |
| *noCIP* | PA14 | 98 | 10 | 11 | 94 | 0 | 0 | 0 | 213 | 118 |
|  |  | 46% | 5% | 5% | 44% | 0% | 0% | 0% |  |  |
|  | **PA14_lys\*** | 162 | 63 | 36 | 74 | 13 | 5 | 31 | 384 | 286 |
|  |  | 42% | 16% | 9% | 19% | 3% | 1% | 8% |  |  |
| | ΔCRISPR | 110 | 27 | 9 | 68 | 0 | 5 | 0 | 219 | 162 |
|  |  | 50% | 12% | 4% | 31% | 0% | 2% | 0% |  |  |
| | ΔCRISPR_lys | 117 | 29 | 20 | 74 | 0 | 3 | 0 | 243 | 153 |
|  |  | 48% | 12% | 8% | 30% | 0% | 1% | 0% |  |  |
| *CIP* | PA14 | 82 | 10 | 14 | 97 | 0 | 1 | 0 | 204 | 121 |
|  |  | 40% | 5% | 7% | 48% | 0% | 0% | 0% |  |  |
|  | **PA14_lys\*** | 150 | 42 | 55 | 90 | 14 | 6 | 41 | 398 | 250 |
|  |  | 38% | 11% | 14% | 23% | 4% | 2% | 10% |  |  |
| | ΔCRISPR | 63 | 15 | 32 | 70 | 0 | 3 | 0 | 183 | 105 |
|  |  | 34% | 8% | 17% | 38% | 0% | 2% | 0% |  |  |
|  | **ΔCRISPR_lys\*** | 121 | 43 | 65 | 64 | 0 | 3 | 30 | 326 | 230 |
|  |  | 37% | 13% | 20% | 20% | 0% | 1% | 9% |  |  |

We identified adaptive mutations as those that change in parallel at >2% frequency in >1 population (Table 1). These loci included non-coding regions like the CRISPR array or intergenic regions, as well as open reading frames. In total, we identified 233 adaptive loci and 1033 mutations. Viral insertions were a minority of both categories (adaptive loci: 2.3%, n = 40). Three loci were mutated in all genotypes and conditions at high frequencies (>25%): *wspA*, *wspF*, and *morA* are crucial regulators of the planktonic-biofilm transition and common targets of selection in both laboratory and infection conditions (*14*, *55*, *56*). Mutations in *wspA* and *wspF* are known to result in elevated c-di-GMP levels and biofilm production, especially a WspA 285-298 aa deletion and frameshift mutations in WspF 154 aa (*55*), both of which were recovered in our populations. Similarly, *morA* encodes a c-di-GMP degrading phosphodiesterase, and mutations lead to an elevated biofilm phenotype (*57*). We tested whether the experimental conditions had a significant impact on both total and adaptive mutations using a PERMANOVA on a presence-absence Jaccard distance matrix of both mutation sets. In both cases, the nested lysogeny:induction term had the strongest effect (all mutations: R^2^ = 6.1%, *P* = 0.0001; adaptive mutations: R^2^ = 5.5%, *P* = 0.0015) (Fig S2E, S2F).

To link adaptive genes to specific selective conditions, we identified adaptive loci (as above) that were present in at least two populations one condition *and* absent in all populations from the other conditions (background: CRISPR vs ΔCRISPR; infection status: lysogen vs no lysogen; environment: CIP vs noCIP; induction: inducing vs noninducing). For example, a variant gene present in two CIP populations and absent in all noCIP populations would be called adaptive in the CIP condition. These variants could be at the same or different nucleotide positions within the same locus, and could include new viral insertion sites as well as host mutations. One gene could be adaptive to multiple conditions. In total, these criteria matched 147 adaptive loci (17% of all variant loci), encompassing 496 adaptive mutations that can be assigned to a specific condition (Fig 2B, Supplementary Data 1). In brief, *lasR* and *rpoB* were the highest-frequency genes specific to the noCIP condition; while *fleQ, dipA, cheA, orfK,* and *pelB* alleles were unique to CIP-evolved populations. *gacA*, a biofilm regulator (*58*) was specific to ΔCRISPR backgrounds evolve in noCIP; similarly, its partner *gacS* was specific to no-induction backgrounds. PA14 lysogen-specific genes included a range of CRISPR-related alleles, including self-targeting spacer mutations and *cas3, cas5, cas7,* and *cas8* variants. PA14_28970, an RNA polymerase sigma factor, and *ptsN*, a nitrogen regulatory protein, were both adaptive in only nonlysogen backgrounds.

### Standing adaptive variation associated with induction by CRISPR self-targeting

Consistent with our previous work (*8*), CRISPR self-targeting lysogens in both the presence and absence of CIP evolved mutations resolving CRISPR self-targeting via both point mutations and transpositional mutations causing polylysogeny. All noCIP-evolved, PA14_lys populations also resolved CRISPR-mediated genetic conflict by inactivating CRISPR through deletion of the locus (strong induction resolution) or self-targeting spacer (weaker induction resolution) or both (Fig S3) (*8*). These unique large CRISPR deletions were never fixed but instead were found at an average of 40% frequency in all PA14_lys populations (frequencies ranged from 20.3% – 59.5%). Similar diversity in CRISPR alleles were also identified in 4 of the 6 CIP-evolved, PA14_lys populations. It was striking that two +CIP PA14_lys populations did not evolve high frequency CRISPR resolution at all although they have lower PFUs and higher OD600s; these two populations, PA14lys_CIP_2 and PA14lys_CIP_4, correspond to the two aforementioned populations with fixed ciprofloxacin resistance mutations. Taken together, these data show that a diversity of adaptive mutations persist as CRISPR targeting lysogens evolve to resolve induction.

### Standing adaptive variation associated with induction by ciprofloxacin

To investigate the genetic basis of evolved resistance in these populations, we specifically assessed mutations which were implicated in ciprofloxacin resistance in the literature (Fig 3). Lysogen populations together had twice as many mutated CIP^R^ genes on average as non-lysogen populations, and had multiple alleles in the same resistance locus per population (Fig 3A). Lysogen populations also accumulated multiple mutations in *mexZ*, *mexK, gyrB* and *orfN* (Fig 3B). Negative regulators like *mexS, nfxB,* and *mexZ* (which repress the MexEF-OprN, MexCD-OprJ, and MexXY-OprM cassettes respectively (*29*, *31*, *59*)) were the most affected: 75% (33/44) of mutations in these genes were small indels, followed by 22% nonsynonymous mutations (10/44), and 2% viral insertion (1/44), consistent with the interruption of negative regulation, leading to increased efflux pump expression. Mutations in *orfN,* a non-canonical ciprofloxacin resistance determinant identified previously in experimental evolution (*56*), only achieved high frequencies in +CIP lysogen populations (Fig S4). Six alleles fixed at 100% frequency; GyrA G75A, MexS A9fs, and a nonsynonymous mutation in the NfxB stop codon all reached fixation in non-lysogens, while only gyrase alleles fixed in lysogen populations (GyrA T83I and A179V, and GyrB S466Y).

**Figure 3.**
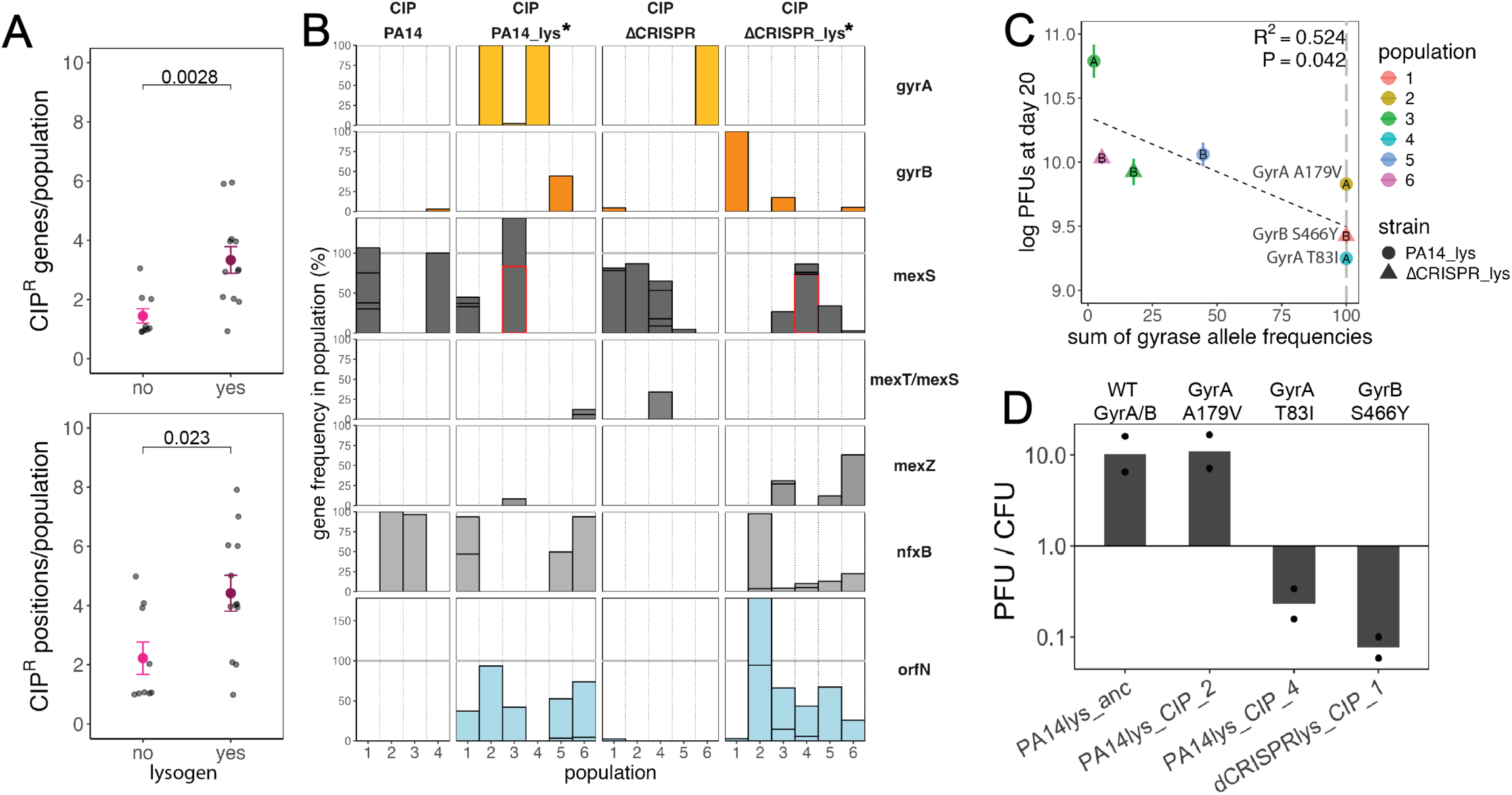
Gyrase allele evolution resolves lysogen induction and reduces AR gene diversity. A) Count of the number of ciprofloxacin resistance genes (top) and resistance alleles (bottom) in non-lysogens and lysogens. B) Bar graph of allele frequency per gene (rows) in each replicate population. Stars next to the labels indicate inducing lysogen backgrounds. Gray line at 100%; note the variable y-axis. Red line denotes if the mutation was caused by viral transposition. One 3% mutation in *oprM* in ΔCRISPRlys_CIP_6 and one 2% mutation in *parC* in ΔCRISPRlys_CIP_3 is not shown, for clarity, but were included in the counts in A). C) Scatter plots of viral titer in each population against the sum of all gyrase alleles in that population. Color represents the population replicate; shape represents the strain. Inscribed letters indicates which gyrase gene was affected. Fixed gyrase alleles are labeled with the exact mutation. D) Relative viral titer of CIP-evolved populations after overnight growth in LB. Bars are the means; points are biological replicates.

Mutations in *gyrA* or *gyrB* inhibit binding of ciprofloxacin to the gyrase to rescue gyrase function (*29*). Of four fixed gyrase mutations – GyrA G75A, T83I and A179V, and GyrB S466Y – only one, GyrA G75A, was in a nonlysogen population (ΔCRISPR_CIP_6) (Fig S5A). Interestingly, populations with fixed gyrase mutations had significantly fewer CIP^R^ alleles in other efflux resistance genes than lysogens without a fixed gyrase allele (Fig S5B), indicating strong selection for these loci and a selective sweep establishing resistance. We tested whether these variant gyrase alleles resolved provirus induction by correlating gyrase allele frequency with the final viral titer of each lysogen population at the end of the experiment. As expected, these variables were negatively correlated (Fig 3C). Viral titer was also persistently lower after overnight growth of the populations in strains with certain gyrase alleles (Fig 3D). In both analyses, the population with a fixed GyrA A179V allele (PA14lys_CIP_2) had viral titers comparable to other populations, while the other mutations *gyrA* and *gyrB (*GyrA T83I and GyrB S466Y) were associated with the decreased viral titers. Both latter mutations occur within the quinolone resistance determining region (QRDR), and the T83I mutation in GyrA is well-known to confer high levels of resistance (*60*). In contrast, viral titer had no relationship with either CIP^R^ gene count, total CIP^R^ gene frequency, or allele frequency of efflux pump alleles (*mexS* and *nfxB*), *orfN, wspF,* or CRISPR self-targeting resolution alleles. This suggests that in lysogens, efflux pump alleles confer intermediate antibiotic resistance that does not stop viral induction. Where *gyrA* or *gyrB* QRDR variants occur, they stop viral induction and are more strongly selected for in lysogens, even under sublethal doses of antibiotic and in the presence of CRISPR self-targeting.

### Induction significantly impacts standing variation in adaptive genes and mutations

Because we found higher numbers of mutations in antibiotic resistance and CRISPR loci in lysogen populations, and lower numbers of mutations in populations which had resolved provirus induction through gyrase evolution, we tested whether viral induction promotes diversification of adaptive evolution across the genome in experimental *Pseudomonas aeruginosa* populations. We were able to differentiate the effect of induction from lysogeny by looking at the noCIP-evolved ΔCRISPR lysogens. Overall induction also had a significant effect on the total number of mutations per population; inducing lysogens had more mutations than both all other populations, and when we compared inducing lysogens to the non-inducing lysogen group (Fig 4D; Kruskal-Wallis test, *P* = 0.001 and *P* = 0.015, respectively). Importantly, this effect remained when we excluded mutations due to viral insertion (Fig 4D; Kruskal-Wallis test of all populations, inducing vs. non-inducing, *P =* 0.001).

**Figure 4.**
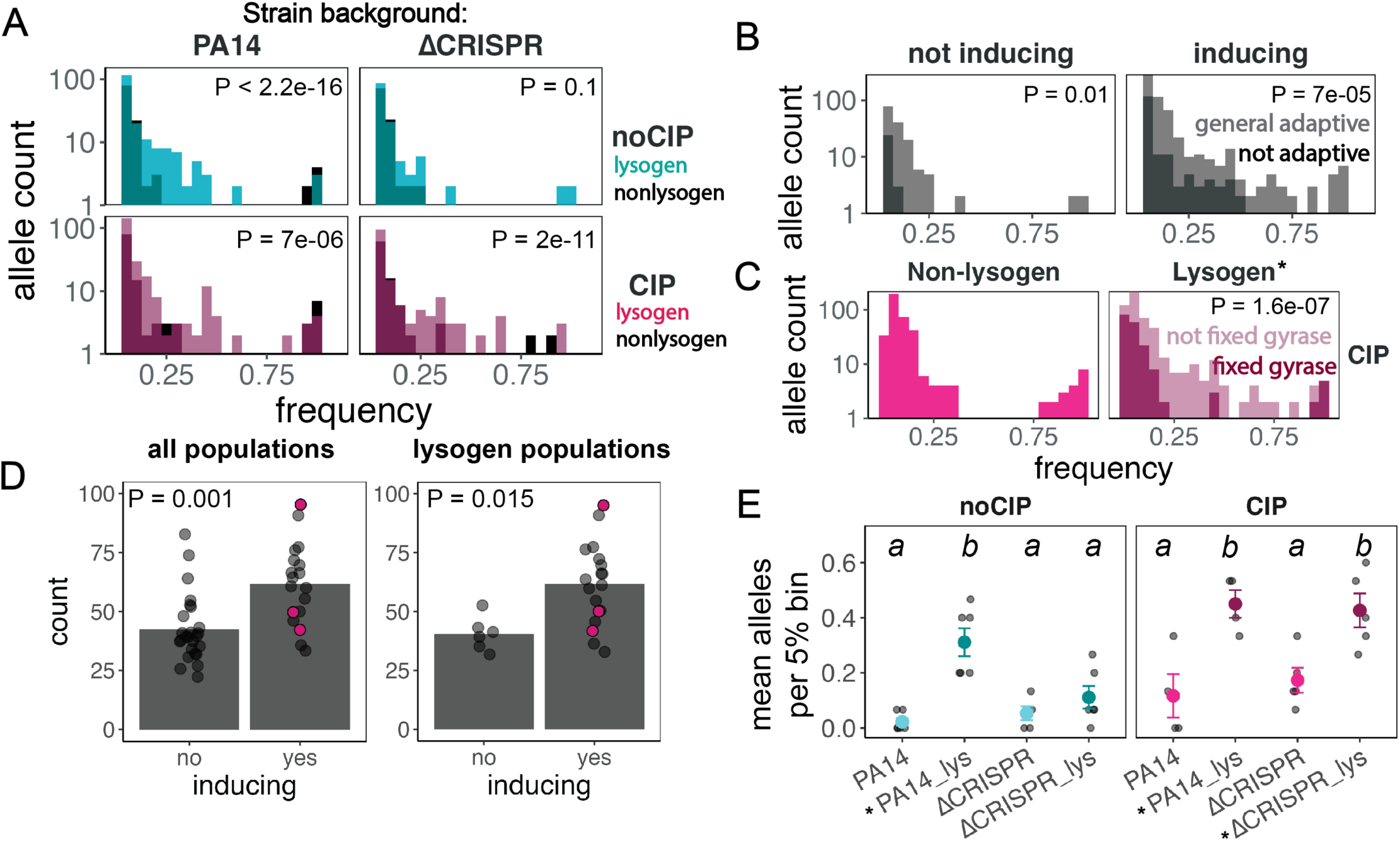
Inducing lysogens have more intermediate frequency alleles than other strains. A) Histogram of allele frequencies of all population in a strain background. Nonlysogens are gray, lysogens are a transparent color. B) Allele frequency histogram of generally adaptive (>2% in >1 population) vs. not adaptive in inducing vs. non inducing strains. C) Allele frequency histogram of nonlysogens and lysogen populations with a fixed gyrase allele (dark pink), and lysogen populations without a fixed gyrase allele (transparent pink). In A-C, significance was tested with a KS test. Inset p-values are the significance of the difference between the two distributions. D) Bar graphs of the counts of all mutations per each population, in all populations (left) and only lysogens (right). Highlighted pink points are lysogen populations with a fixed gyrase mutation (the highest one is in, PA14lys_CIP_2 with a gyrase allele which does not resolve induction). Significance was tested with a Kruskal-Wallis test. E) The mean number of alleles per 5% bin of histograms of all allele frequencies from 25-95%. Letters indicate the results of an ANOVA, with Tukey post-hoc test for individual strains across CIP treatments. Categories with overlapping letters are not significant. Throughout, stars denote inducing lysogen strains.

Inducing lysogen strains had a significantly different allele distribution from non-inducing strains. Individual comparisons between lysogens and nonlysogens showed that in every case, allele frequencies in inducing lysogens were significantly skewed toward higher intermediate frequencies compared to the nonlysogen (Fig 4A); differences were not significant between the noninducing lysogen and nonlysogen populations (noCIP ΔCRISPR backgrounds) (Fig 4A). In both the presence and absence of ciprofloxacin, inducing populations had significantly different distributions which featured a prominent increase in intermediate frequencies increasing overall diversity (Fig 4B). This shifted frequency spectra however, depended on the absence of a fixed gyrase allele: the CIP-evolved lysogens with a fixed gyrase allele had fewer alleles at intermediate frequencies than lysogens which hadn’t effectively resolved induction (Fig 4C). The shifted distribution of inducing lysogens (not including the three fixed-gyrase lysogen populations) could be explained either by having more intermediate mutations overall, or more alleles in a particular frequency bin (which would suggest linkage between those alleles).

We compared the number of alleles per 5% bin of frequencies in the 25-95% range, and found that inducing lysogens also had significantly more alleles per bin than their nonlysogen, non-inducing counterpart (Fig 4E). This difference resulted primarily from a higher number of intermediate alleles as a whole, rather than more coexisting mutations in the same bin, even when we increased frequency bin size to 10%. These intermediate frequencies could also represent stably maintained subpopulations (soft sweeps, where multiple alleles in the same gene persist at intermediate frequencies through clonal competition, but the gene itself is mutated in 100% of the population (*61*)).

## Discussion

Here, we show that viral induction promotes diversifying adaptive evolution in lysogen populations. This diversification was reflected in the increased absolute number of adaptive mutations and in the number of sequential and coexisting adaptive mutation that are found at intermediate frequencies in the population’s metagenome. As a result, in the presence of ciprofloxacin, we found that lysogen populations maintained twice as many antibiotic resistance alleles as nonlysogens.

Why does lysogen induction diversify adaptive mutation? We propose that in a clonal population, lysogen induction shifts the population mutation-selection regime relative to non-inducing populations. Desai and Fisher describe four regimes resulting from a continuum of weak and strong selection balanced by weak and strong mutation in populations of different sizes (*62*). Along this continuum the selective regime changes from one that results in successional selective sweeps, where populations have low variation at any one time point, to one of concurrent mutation where clonal competition among many mutations of similar effect coexist leading to increased variation across time. In this study, we demonstrate that GxGxE interactions move populations across this continuum caused by virus induction. In the CRISPR-targeting background there are both weak and strong effect alleles (spacer vs large CRISPR deletions). In this context, clonal interference among CRISPR deletions, occurring frequently through transposition, maintains diversity, and smaller effect mutations (spacers) coexist by decreasing the selection differential if they occur early by chance. The sublethal ciprofloxacin condition changes the ruggedness of this landscape by increasing the number of small effect mutations (efflux pumps through mutation and transposition) and adding large effect mutations in the gyrase genes. Only the lucky few evolve a large effect gyrase mutation which removes diversity through a selective sweep. Importantly, all three fixed gyrase mutations in lysogen populations were in the quinolone resistance determining regions of *gyrA* and *gyrB* (*60*), and the frequency of gyrase variant alleles negatively correlated with viral titer and the number of adaptive mutations within populations that contain them. Additionally in some populations, the clonal interference from weaker effect mutations maintains diversity and promotes sequential mutations to occur in persistent competing lineages. The effect of classical efflux pump mutations in *mexS* and *nfxB* is dampened, which allows the selection and maintenance of other, less effective resistance alleles like *orfN,* which readily arises in laboratory evolution in a range of antibiotic treatment conditions (*56*, *63–67*).

Our findings are consistent with a model where increased mutation rates due to viral transposition contribute a larger number of beneficial alleles of equal benefit (CRISPR deletions, efflux pumps) which may evolve in the background of a sweeping beneficial allele or persist through clonal interference depending on when they evolve in the course of the changing selective landscape (*62*). Both the standing variation and the rate of beneficial mutation accumulation increase logarithmically with population size and/or mutation rate (*62*). Viral induction increases mutation through transposition but also decreases population size which should counteract this effect. It is possible that increases in population size due to reduction in viral lysis, combined with the increase in mutations of small effect, together play a role in preserving the increased adaptive variation we observe by the end of the experiment. Overall, the combination of increased mutations, a more rugged adaptive landscape, and structural variations shift the mutation-selection balance and result in evolutionary trajectories that are diverged between lysogens and nonlysogens.

This work has several limitations. First, we only performed sequencing at one time-point, limiting our ability to understand relative rates of evolution in lysogens and non-lysogens and how pervasive these differences are throughout the course of the experiment. Second, the timeline was short; ciprofloxacin-treated lysogen populations never resolved their high viral induction rates to baseline levels, which suggests that viral induction would continue to affect the evolution of those populations. Third, because we used a transposable virus with an antibiotic that causes induction, we cannot fully disentangle the relative impact of increased mutation, changes in population size and the selective landscape for antibiotic resistance evolution in general. Future studies with non-transposable but inducible viruses could help to resolve the relative contribution of lysis and mutation to establishing this mutation-selection regime more generally. Fourth, we did not explore heterogeneity in time and space at the level of ciprofloxacin exposure that is likely to exist within treated lungs of CF patients, and impact this evolutionary process. Although ciprofloxacin is used in the treatment of bacterial infections in people with CF, it is often used in combination with tobramycin or other antibiotics (*25*, *68*). Non-DNA damaging antibiotics may still effect provirus induction, and the effect of quinolones on virus induction may also depend on the identity of the virus (*69*).

We find that lysogens evolved greater variation in adaptive alleles under inducing conditions. Lysogen populations overall had more mutations under inducing conditions, whether the induction was due to ciprofloxacin treatment or CRISPR self-targeting. We conclude that viral induction increases adaptive antibiotic resistance gene diversity in lysogens by altering the selective landscape, resulting in more adaptive and more diverse resistance alleles at intermediate frequencies. Many viruses are induced via DNA damage, making this a relationship which is potentially applicable to a wide variety of virus-host pairs. We suggest that if similar trajectories occur *in vivo* in the CF lung, the ancestral strain genotype and its differential evolutionary regimes should be considered in combination before treatment with antibiotics that use similar efflux pumps or through phage therapy. The diversification in efflux pump genes we observe here is likely to complicate phage therapy strategies that target efflux pump receptors (*70*, *71*). The outcome of the multiscale, co-evolutionary dynamic is an important topic for further study in this and other chronic infectious diseases.

## Acknowledgements

We gratefully acknowledge Alvaro Hernandez and Chris Wright of the Roy J. Carver Biotechnology Center at the University of Illinois Urbana-Champaign for sequencing, and Jaya Chandrashekhar for sequencing conversations and help during the evolution experiment. We thank George O’Toole for the strains, and Nkrumah Grant for helpful conversations. We thank Sierra Raglin for advice on the redundancy analysis; Jiayue Yang, Thomas Cowell, and Caroline Oldstone-Jackson for critical reading of early drafts of the manuscript; and the rest of the Whitaker lab for thoughtful discussions. We gratefully acknowledge funding and the interdisciplinary educational framework provided by NSF BII Award 2022049 through the GEMS (Genomics and Eco-Evolution of Multi-Scale Symbioses) Institute, as well as funding from the Allen Foundation Distinguished Investigator Award (RJW) and the Cystic Fibrosis Foundation.

## Supplementary Figures

**Supplementary Figure 1.**
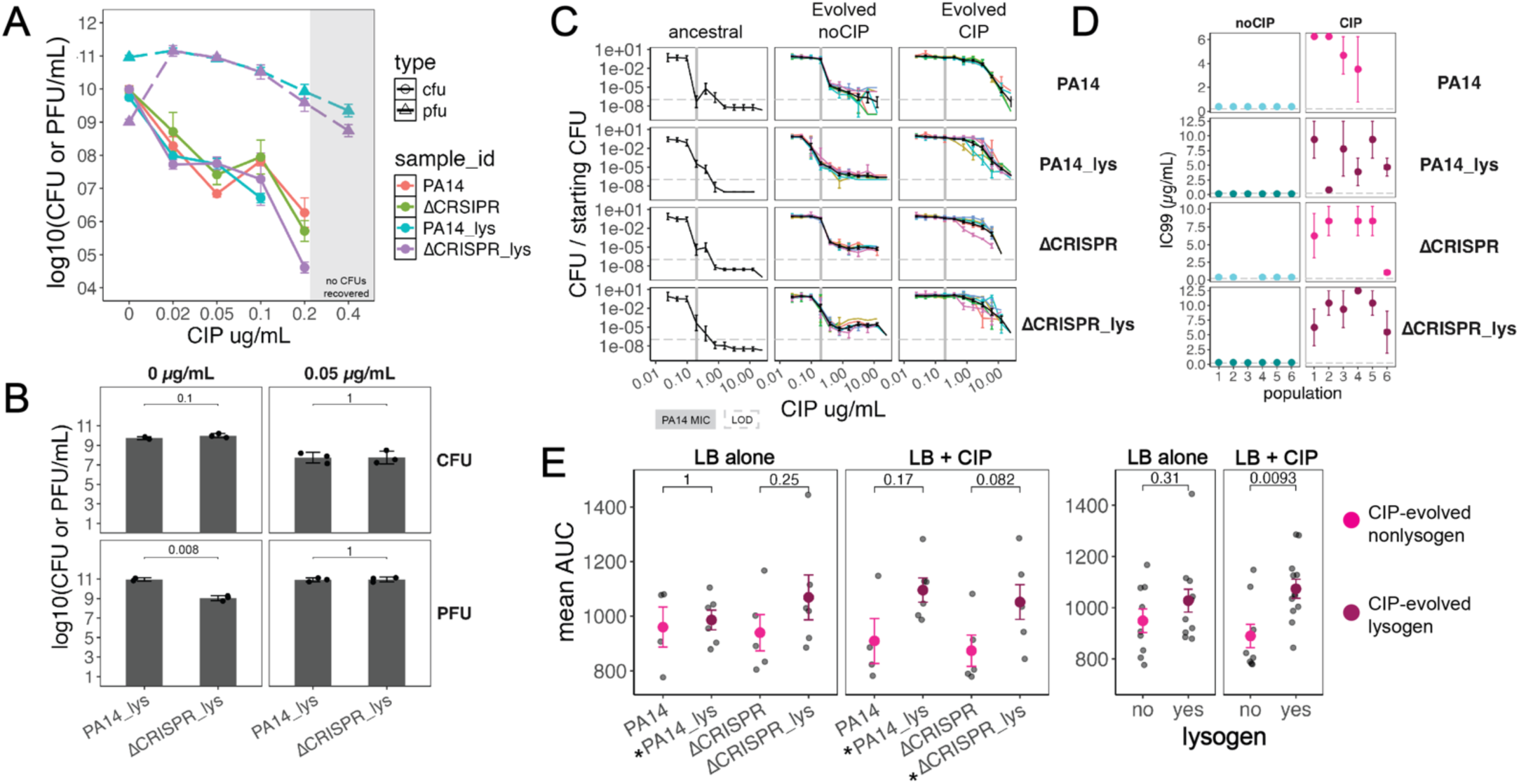
Antibiotic resistance in cultures evolved in 0.05 µg/mL ciprofloxacin. A) Log10 CFU and PFU/mL from ancestral cultures incubated with the indicated CIP concentration overnight. No CFUs were recovered at concentrations greater than 0.2 ug/mL. B) Ciprofloxacin induction induces ΔCRISPR_lys populations to the level of CRISPR-containing PA14_lys populations. C) Death curves of ancestral (left column), noCIP-evolved populations (middle) and CIP-evolved populations (right), grown in LB overnight and serially diluted onto solid media with increasing concentrations of ciprofloxacin. Black lines are the average; error bars are standard error; colorful lines in the background are individual replicates. The gray line is the MIC of ancestral PA14; the gray dashed line is the limit of detection (LOD). D) IC99s of strains evolved in the absence and presence of ciprofloxacin. Error bars are standard error. Points are the means. Dashed line indicates 0.2 µg/mL ciprofloxacin (PA14 MIC). E) The mean area under the curve (AUC) of growth curves of CIP-evolved populations in the presence (right) and absence (left) of CIP, of individual strains or grouped lysogens. Black points are individual replicates; colorful points are the means; error bars are standard error. Significance was tested with Kruskal-Wallis tests. Stars indicate inducing lysogens.

**Supplementary Figure 2.**
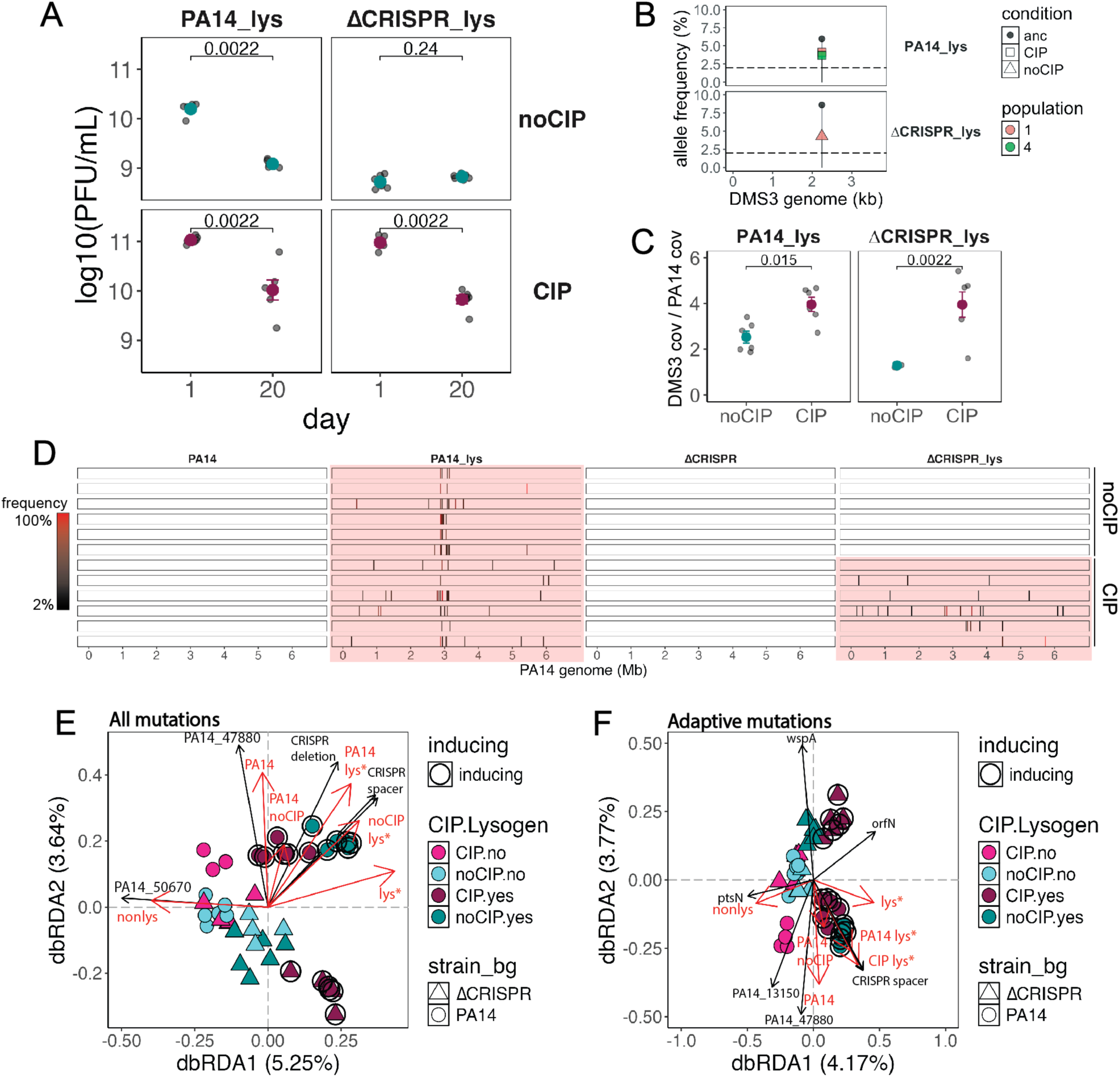
Sublethal ciprofloxacin treatment mobilizes viral transposition around the genome. A) Viral titers at Day 20 in all lysogen populations in log10(PFU/mL). B) Frequency of mutations in the DMS3 genome does not exceed variation in an overnight culture of the ancestor (black dot, “anc”). C) Relative coverage of the DMS3 genome, compared between the same strain, evolved with and without ciprofloxacin. D) Location of new viral insertion sites ≥2% frequency across the genome. Ancestral location is not shown. Color of the bar is the frequency; highlighted populations are inducing lysogens. E, F) Distance-based redundancy analysis plot of Jaccard dissimilarity of E) all mutations and F) adaptive mutations (present in >1 population) in each population. Red arrows and text indicate the main terms of the model; black arrows and text are the gene names of some significant mutations from the envfit function. Arrow length was shortened for visualization. Asterisks indicate induction in the model term, and inducing populations are inscribed with a larger black circle. For both E and F), significance was determined with Adonis2 PERMANOVA. E) Significance of treatments in differentiating all mutation content in populations: Ciprofloxacin, R^2^ = 2.6%, *P* = 0.008; strain, R^2^ = 2.5%, *P* = 0.12; strain x ciprofloxacin, R^2^ = 2.3 %, *P* = 0.568; lysogen:inducing, R^2^ = 6.1%, *P* = 0.001; strain x lysogen:inducing, R^2^ = 2.2%, *P* = 0.553; cipro x lysogen:inducing, R^2^ = 2.2%, *P* = 0.817. F) Significance of treatments in differentiating adaptive mutation content in populations: Ciprofloxacin, R^2^ = 2.6%, *P* = 0.114; strain, R^2^ = 2.6%, *P* = 0.085; strain x ciprofloxacin, R^2^ = 2.3%, *P* = 0.698; lysogen:inducing, R^2^ = 5.5%, *P* = 0.0031; strain x lysogen:inducing, R^2^ = 2.2%, *P* = 0.653; cipro x lysogen:inducing, R^2^ = 2.4%, *P* = 0.324.

**Supplementary Figure 3.**
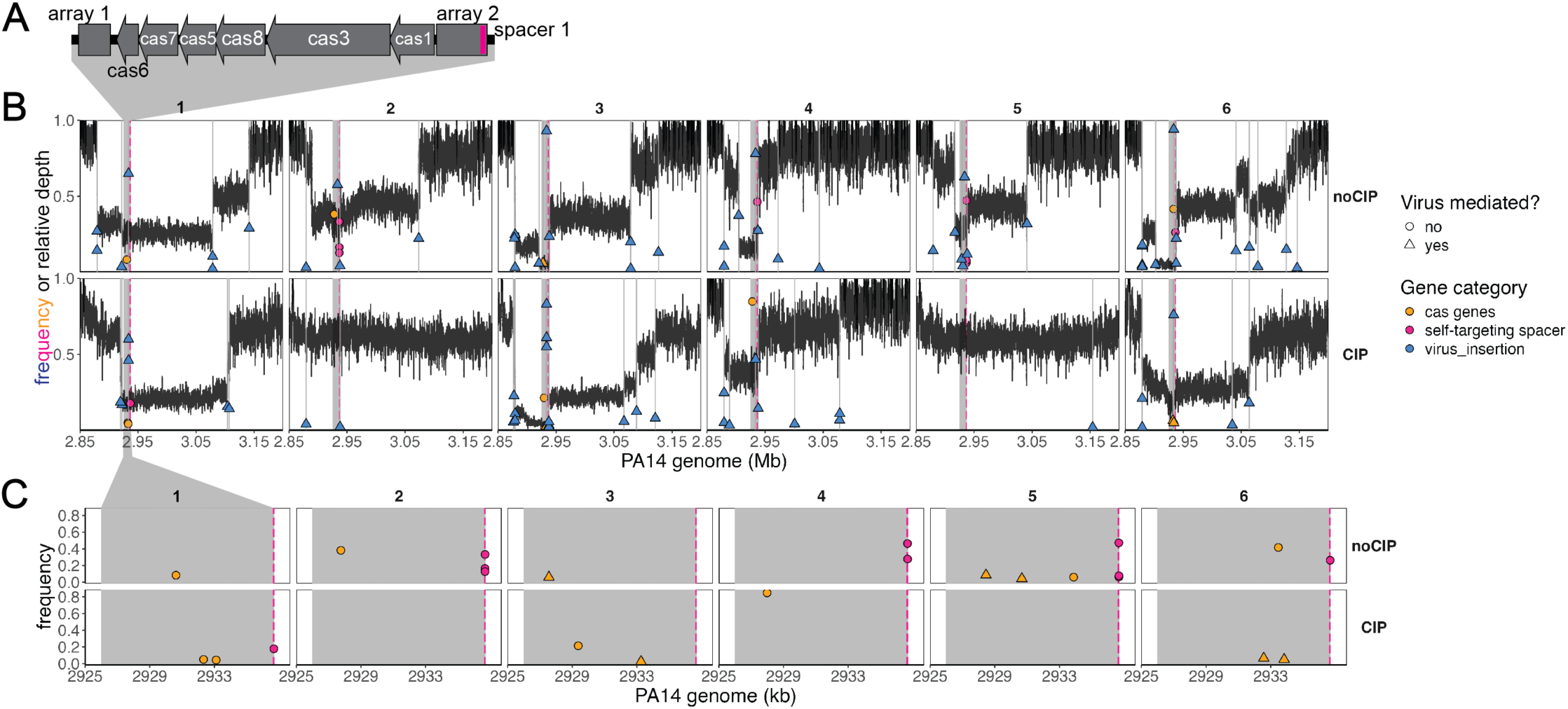
Incomplete and diverse resolution of CRISPR self-targeting in inducing conditions. A) CRISPR arrays in PA14. The self-targeting spacer is highlighted in pink. B) Wide view of the region surrounding the CRISPR locus. Relative depth (depth normalized by mean coverage) is plotted as a black line. Mutations, including viral insertions, are overlaid. Color represents the mutation category. C) Zoomed view of the CRISPR locus. Gray box represents the CRISPR boundaries; color represents category of CRISPR mutation. Dashed pink line indicates the location of the self-targeting spacer.

**Supplementary Figure 4.**
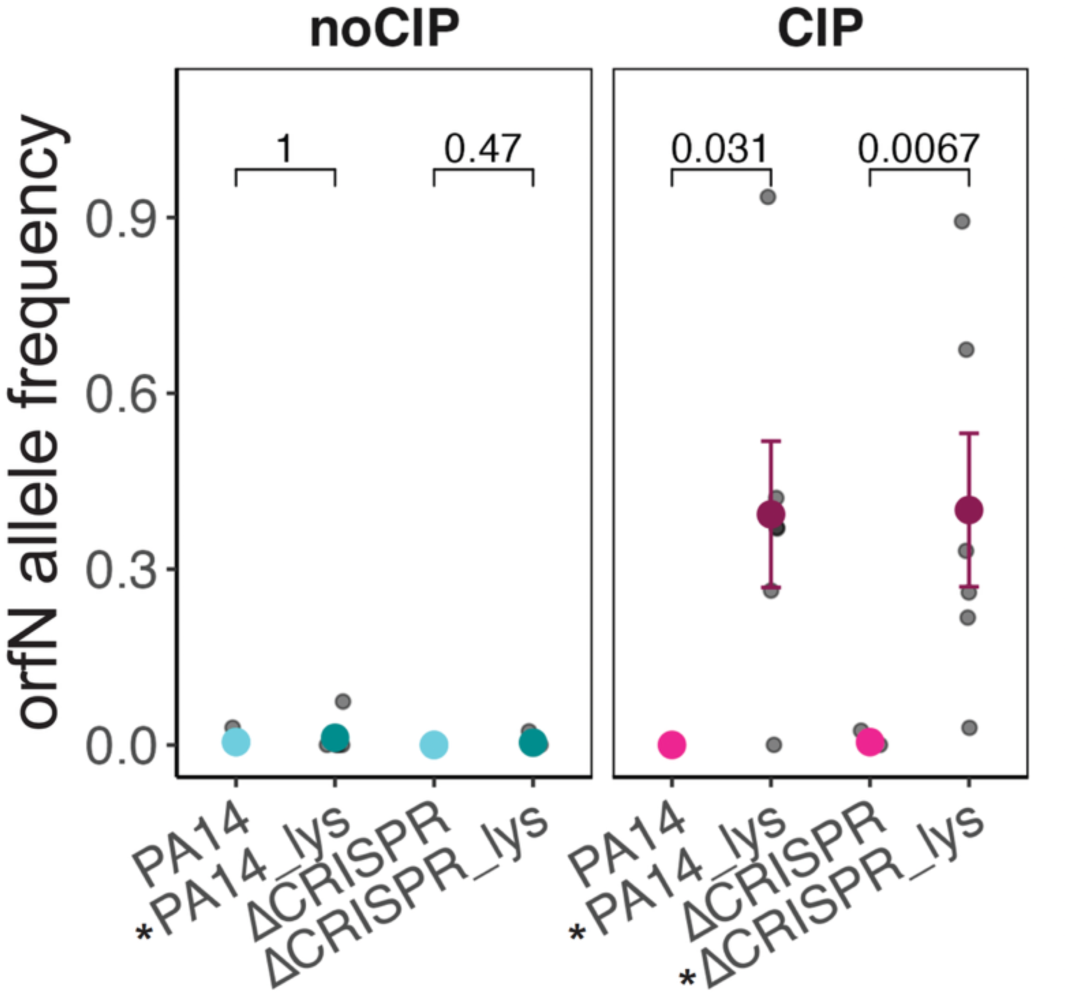
Naturally variable locus is increased in frequence in CIP-evolved lysogens. Frequency of *orfN* gene variants across all populations. In all, points are a population; colored points are the mean; error bars are standard error. Stars indicate inducing lysogen strains.

**Supplementary Figure 5.**
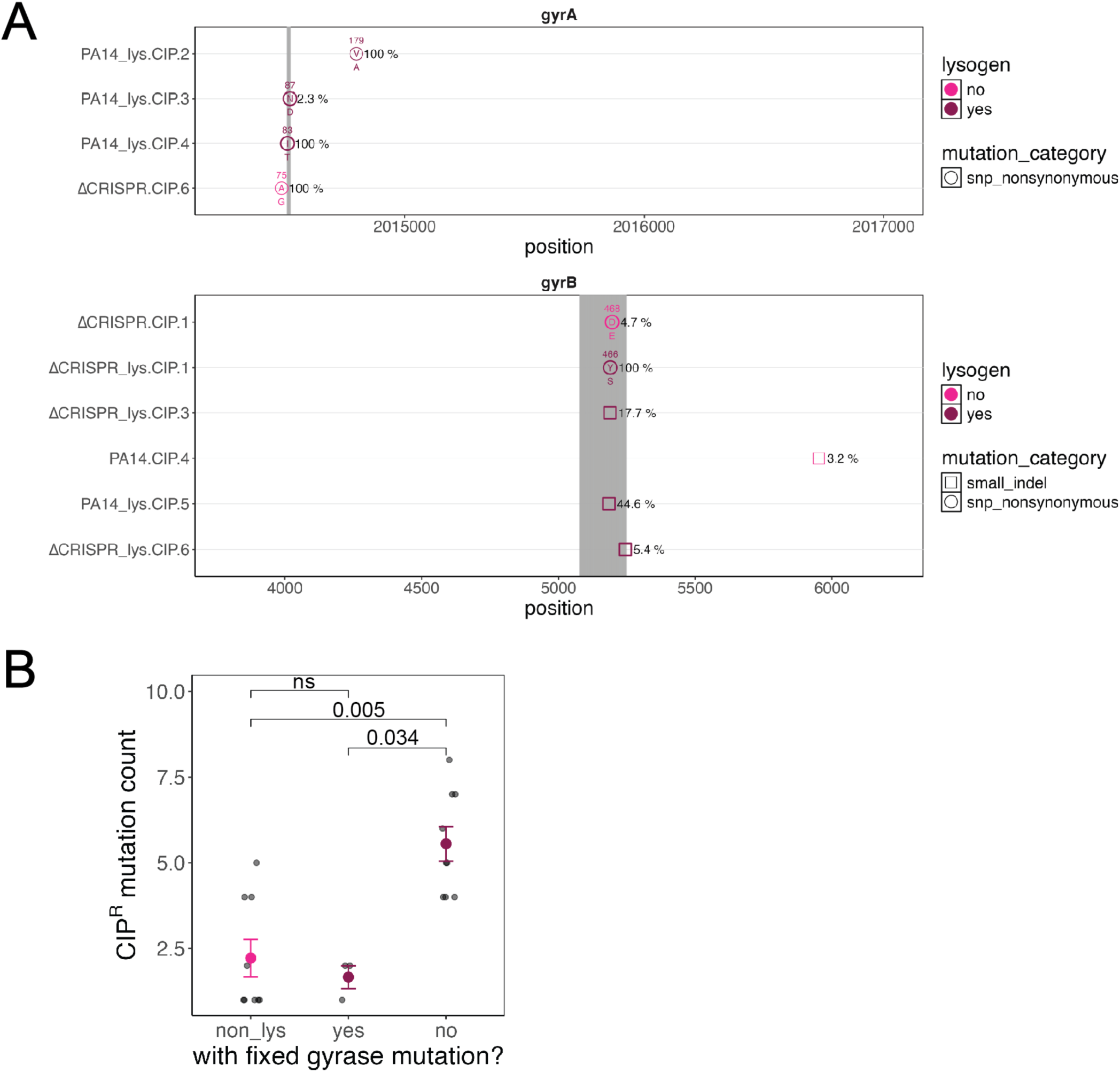
High efficacy gyrase alleles correlate with lower viral induction. A) Map of gyrase allele location in *gyrA* and *gyrB* genes. QRDR (quinolone resistance determining region) is highlighted in gray. X-axis is the position on the gene. Y axis is the population replicate (all were evolved in ciprofloxacin). Shape of the point is the type of mutation. Color represents whether the population is a lysogen. Points are annotated with the variant allele in the center, and the wildtype allele at the bottom of the point. The location of the amino acid is at the top. Frequency is at the side. B) Count of resistance mutations in non-lysogens compared to lysogens with and without a fixed gyrase allele.

